# Mechanism of ASF1 Inhibition by CDAN1

**DOI:** 10.1101/2024.08.08.607204

**Authors:** Samantha F. Sedor, Sichen Shao

## Abstract

Codanin-1 (CDAN1) is an essential and ubiquitous protein named after congenital dyserythropoietic anemia type I (CDA-I), an autosomal recessive disease that manifests from mutations in the *CDAN1* or *CDIN1* (<u>CD</u>AN1 interacting <u>n</u>uclease 1) gene. CDAN1 interacts with CDIN1 and the paralogous histone H3-H4 chaperones ASF1A (<u>A</u>nti-<u>S</u>ilencing <u>F</u>unction 1A) and ASF1B, but its function remains unclear. Here, we biochemically and structurally analyze CDAN1 complexes. We find that CDAN1 dimerizes and assembles into cytosolic complexes with CDIN1 and multiple copies of ASF1A/B. Single-particle cryogenic electron microscopy (cryo-EM) structures of CDAN1 complexes identify interactions with ASF1 mediated by two CDAN1 B-domains commonly found in ASF1 binding partners and two helices that mimic histone H3 binding. We additionally observe that one CDAN1 can recruit two ASF1 molecules and that ASF1A and ASF1B have different requirements for CDAN1 engagement. Our findings explain how CDAN1 sequesters and inhibits the chaperone function of ASF1A/B and provide new molecular-level insights into this enigmatic complex.

## Introduction

Congenital dyserythropoietic anemias (CDAs) are inherited disorders primarily characterized by failure of the erythroid lineage to effectively differentiate into red blood cells. CDAs are classified into subtypes based on distinctive bone marrow morphologies arising from mutations in specific genes^1–4^. Some CDA-associated genes, such as *GATA1*^5^ and *KLF1*^6^, encode transcription factors that primarily function in hematopoiesis. Other CDA genes are linked to fundamental cellular functions. These include *SEC23B*^7,8^, which encodes a component of the COPII coat that mediates anterograde vesicular transport, and *KIF23*^9^ and *RACGAP1*^10^, which encode subunits of the centralspindlin complex required for cytokinesis.

Most genes associated with CDAs are well studied. However, *CDAN1* and *CDIN1* (*C15orf41*), the two genes linked to CDA subtype 1 (CDA-I), encode a protein complex of unknown function^11–14^. A striking feature of CDA-I is the formation of spongy or “Swiss cheese” heterochromatin in erythroblast nuclei, a defect in chromatin compaction not seen in other CDAs^15–20^. Consistent with the possibility that this complex regulates chromatin condensation, CDAN1 is reported to interact with ASF1A and ASF1B^21^, paralogous histone chaperones that we will refer to as ASF1 in interchangeable contexts throughout this study. ASF1 binds the H3-H4 dimer and shuttles between the cytosol and nucleus to facilitate downstream nucleosome assembly^22–27^. CDAN1 is proposed to inhibit this function by sequestering ASF1 in complex with H3-H4 in the cytosol via a ‘B’-domain^21^, a short linear motif containing two consecutive basic residues that is used by multiple ASF1 binding partners^28–32^. CDIN1 contains a predicted PD-(D/E)XK nuclease domain^11^ but no nucleic acid substrates have been conclusively linked to its putative enzymatic activity.

Despite their strong link to CDA-I, CDAN1 and CDIN1 probably mediate a fundamental cellular function that is not restricted to erythropoiesis. Both *CDAN1* and *CDIN1* are ubiquitously expressed and essential^33–35^. Notably, knocking out either gene is embryonic lethal in mice prior to the onset of red blood cell production^33,34^. In addition, basic questions regarding the molecular interactions of these proteins remain outstanding. In this study, we integrate genetic, biochemical, and structural approaches to investigate the interactions of CDAN1 with its binding partners. We endogenously tag CDAN1 and CDIN1 to show that these proteins form an obligate cytosolic complex that simultaneously binds ASF1 but not histones. Biophysical measurements and single-particle cryogenic electron microscopy (cryo-EM) additionally reveal that CDAN1 dimerizes and can engage multiple ASF1 molecules through distinct B-domains and helices that mimic histone H3 binding to ASF1. Furthermore, we find that these elements engage ASF1A and ASF1B differently. These results elucidate the interactions used by CDAN1 to recruit and inhibit the chaperone function of ASF1, thus providing new mechanistic insights into this essential complex.

## Results

### Endogenous CDAN1 and CDIN1 assemble together with ASF1

To investigate CDAN1 and CDIN1 in dividing cells without potential artifacts of overexpression, we used CRISPR-Cas9 methods to endogenously tag CDAN1 or CDIN1 in Flp-In 293 T-REx cells with a C-terminal HaloTag-FLAG (HF) epitope (**Supplemental Fig. 1a**). CDAN1 is reported to interact with CDIN1 and ASF1 through distinct binding sites^21,36,37^, but it is unclear if CDAN1 can bind these factors at the same time. Consistent with prior studies, immunoprecipitations of endogenous CDAN1-HF recovered CDIN1 and both ASF1A and ASF1B (**Fig. 1a**, lane 9). Immunoprecipitations of endogenous CDIN1-HF also recovered CDAN1, as well as ASF1A and ASF1B (**Fig. 1a**, lane 8, **and Supplemental Fig. 1b**). In addition, endogenous CDIN1 was significantly depleted from the flow-through of CDAN1-HF immunoprecipitations (**Fig. 1a**, lane 6) and vice versa (**Fig. 1a**, lane 5), indicating that CDAN1 and CDIN1 primarily exist in complex with each other. These findings suggest that CDAN1 can simultaneously bind CDIN1 and ASF1 and confirm that the HF tag does not disrupt the interactions between these factors.

**Figure 1.**
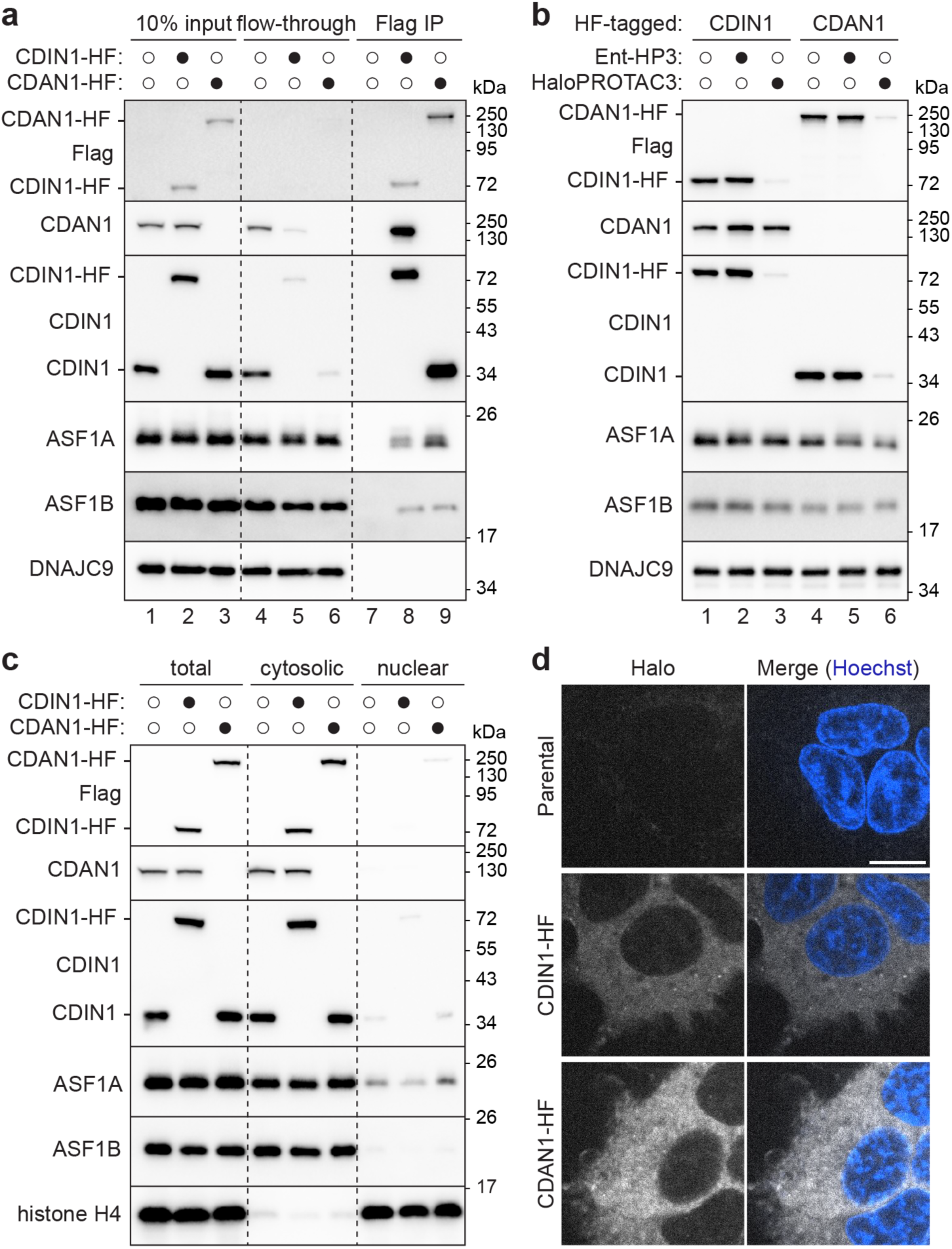
CDAN1 and CDIN1 assemble together with ASF1 in the cytosol. **a,** Endogenous CDAN1 and CDIN1 assemble with ASF1. Flp-In 293 T-REx cells without or with a C-terminal HaloTag-FLAG (HF) tag on endogenous CDIN1 or CDAN1 were lysed (input), subjected to anti-FLAG immunoprecipitations (IP), and analyzed by SDS-PAGE and immunoblotting; representative of 3 independent replicates. Note: the CDAN1 antibody raised against a C-terminal epitope does not recognize CDAN1-HF. Where applicable, different protein populations recognized by the same antibody are labeled. **b,** CDIN1 stability requires CDAN1. Parental, CDIN1-HF, or CDAN1-HF cells were treated without or with 500 nM HaloPROTAC3 (HP3) or an inactive enantiomer (ent-HP3) for 72 hr, lysed, and analyzed by SDS-PAGE and immunoblotting; representative of at least 3 independent replicates. **c,** CDAN1 and CDIN1 are cytosolic. Parental, CDIN1-HF, or CDAN1-HF cell lysates (total) were fractionated to separate cytosolic and nuclear contents and analyzed by SDS-PAGE and immunoblotting; representative of 3 independent replicates. **d,** Representative live-cell images of parental, CDIN1-HF, and CDAN1-HF cells labeled with JFX650 HaloTag ligand and Hoechst. Scale bar, 10 µm.

Next, we used HaloPROTAC3 (HP3), a small molecule degrader for HaloTag fusion proteins^38,39^, to acutely deplete CDAN1-HF or CDIN1-HF (**Fig. 1b**). We achieved robust and specific degradation of each HF-tagged protein upon treatment with HP3 (**Fig. 1b**, lanes 3 and 6) but not an inactive isomer (Ent-HP3) (**Fig. 1b**, lanes 2 and 5). Importantly, acute degradation of CDAN1-HF destabilized endogenous CDIN1 (**Fig. 1b**, lane 6). In contrast, degradation of CDIN1-HF did not change endogenous CDAN1 levels (**Fig. 1b**, lane 3). We observed the same dependency of CDIN1 levels on CDAN1 using siRNA-mediated knockdowns^19^ (**Supplemental Fig. 1c**), while ASF1A and ASF1B levels were not impacted by depletion of either protein. Thus, the stability of CDIN1 depends on CDAN1^36^, supporting the interpretation that CDIN1 forms an obligate complex with CDAN1.

The interaction between CDAN1 and ASF1A is cytosolic^21^. However, several studies report nuclear localization of CDIN1 and/or CDAN1^19,20,36,37,40–42^. Given our results that CDAN1 can form a complex with both ASF1 and CDIN1 and that most of CDIN1 appears to be bound to CDAN1, we examined the localization of these proteins using independent methods. Using our endogenously tagged cell lines, we performed biochemical fractionations followed by immunoblotting with validated antibodies (**Fig. 1c**), as well as live imaging after labeling the HaloTag fusion proteins with a fluorophore-conjugated ligand (**Fig. 1d**). Both approaches demonstrate that endogenous CDAN1 and CDIN1 are primarily cytosolic in Flp-In 293 T-REx cells (**Fig. 1c,d**). This result was initially unexpected considering prior immunofluorescence results using CDIN1 antibodies^19,20,40^. We confirmed that for the antibody previously used in HEK293 cells^40^, immunofluorescence signal did not decrease upon siRNA-mediated knockdown of CDIN1 (**Supplemental Fig. 1c,d**) or HP3-mediated degradation of CDIN1-HF (**Supplemental Fig. 1e,f**), suggesting that the antibody is nonspecific^37^. Our data therefore show that endogenous CDAN1 assembles into a cytosolic complex with CDIN1 and ASF1 paralogs.

### CDAN1 complexes contain multiple copies of each subunit

Next, using transiently transfected Expi293 cells, we performed tandem purifications of Strep-tagged CDAN1 (ST-CDAN1) in complex with FLAG-tagged CDIN1 (F-CDIN1) and HA-tagged ASF1A (**Fig. 2a**, C:C:A complex for CDAN1:CDIN1:ASF1A), or with only FLAG-tagged ASF1A (F-ASF1A) (**Fig. 2b**, C:A complex for CDAN1:ASF1A). Both purifications isolated complexes with approximately stoichiometric amounts of each recombinantly expressed component.

**Figure 2.**
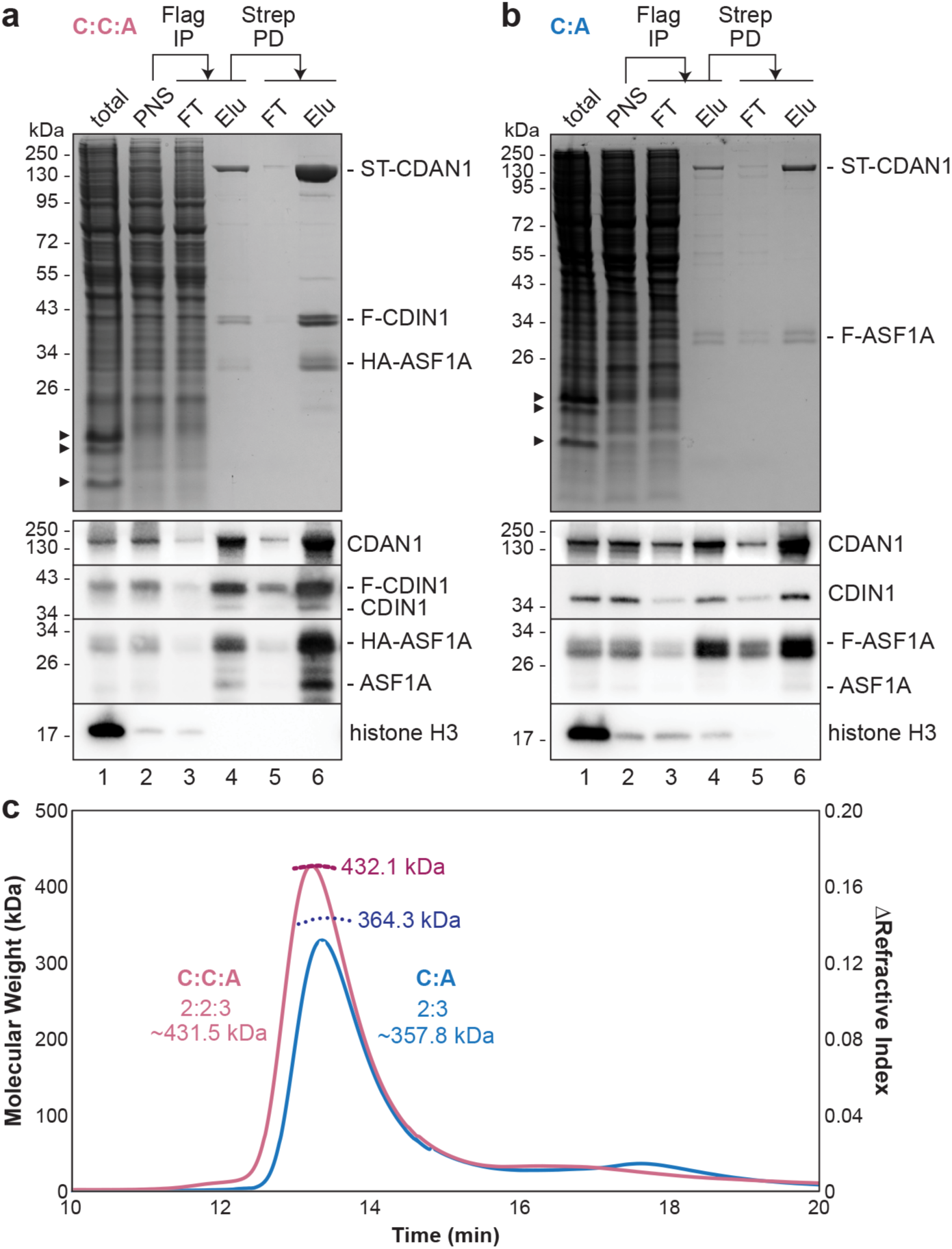
CDAN1 complexes contain multiple copies of each subunit. **a,** Strep-tagged CDAN1 (ST-CDAN1), FLAG-tagged CDIN1 (F-CDIN1), and HA-tagged ASF1A were overexpressed in Expi293 cells by transient transfection for 72 hr. The cells were then lysed, and the post-nuclear supernatants (PNS) were subjected to anti-FLAG immunoprecipitation (FLAG IP) followed by Strep-Tactin pulldowns (Strep PD) to purify the CDAN1-CDIN1-ASF1A (C:C:A) complex. The input, flow-through (FT), and elution (Elu.) samples were analyzed by SDS-PAGE and Coomassie staining (top) or immunoblotting (bottom); representative of at least 2 independent replicates. Arrowheads denote histones. **b,** As in **a**, except with overexpression of ST-CDAN1 and FLAG-tagged ASF1A (F-ASF1A) to purify the CDAN1-ASF1A (C:A) complex. **c,** Size exclusion chromatography with multi-angle light scattering (SEC-MALS) of the C:C:A and C:A complexes determined molecular weights consistent with 2 copies of CDAN1, 3 copies of ASF1A, and, in the C:C:A complex, 2 copies CDIN1. The predicted molecular weights of these stoichiometries are indicated.

Immunoblotting of samples collected at different steps of the C:C:A and C:A purification procedures revealed several notable insights. First, immunoprecipitation of F-CDIN1 substantially depleted ST-CDAN1 from the lysate (**Fig. 2a**, compare lanes 2 and 3), indicating that these two proteins are usually associated with each other, even when overexpressed. Second, both purifications contained low but detectable amounts of endogenous CDIN1 and ASF1A, even if their tagged counterpart was used for purification (**Fig. 2a,b**, lanes 4 and 6). This observation suggests that multiple copies of each factor are present in the complexes, discussed below. Finally, immunoprecipitation of F-ASF1A recovered a detectable level of histone H3 (**Fig. 2b**, lane 4) that was lost after subsequently pulling down on ST-CDAN1 (**Fig. 2b**, lane 6). H3 also was not observed after F-CDIN1 immunoprecipitation (**Fig. 2a**, lane 4). This suggests that CDAN1 complexes do not sequester the histone H3-H4 dimer together with ASF1 in the cytosol, as previously suggested^21^.

The C:C:A and C:A complexes migrated as single peaks by size exclusion chromatography (SEC) (**Fig. 2c and Supplemental Fig. 2a**), indicating stable assembly of the subunits, and negative stain EM images of the C:C:A complex showed discrete particles (**Supplemental Fig. 2b**). Both the C:C:A and the C:A complexes also migrated at a larger size than expected for an assembly containing one copy of each subunit^37^. To determine the molecular weights of these complexes more precisely, we performed SEC with multi-angle light scattering (SEC-MALS) and found that the C:A complex likely contains two copies of CDAN1 and three copies of ASF1A (**Fig. 2c**). Similarly, the measured molecular weight of the C:C:A complex is most consistent with a stoichiometry of two CDAN1, two CDIN1, and three ASF1A. These data suggest that CDAN1 dimerizes and is capable of binding multiple ASF1A molecules, despite containing only one described B-domain^21^.

### Cryo-EM structures of CDAN1 complexes

To understand the interactions that mediate CDAN1 dimerization and ASF1A association, we determined single-particle cryo-EM structures of the C:C:A complex (**Fig. 3, Supplemental Fig. 3 and 4, and Table 1**). 3D classification strategies allowed us to obtain cryo-EM maps at overall resolutions ranging from 3.0 to 3.5 Å of a CDAN1 dimer with either two or three ASF1A molecules clearly resolved (**Fig. 3b,c and Supplemental Fig. 3 and 4**).

**Figure 3.**
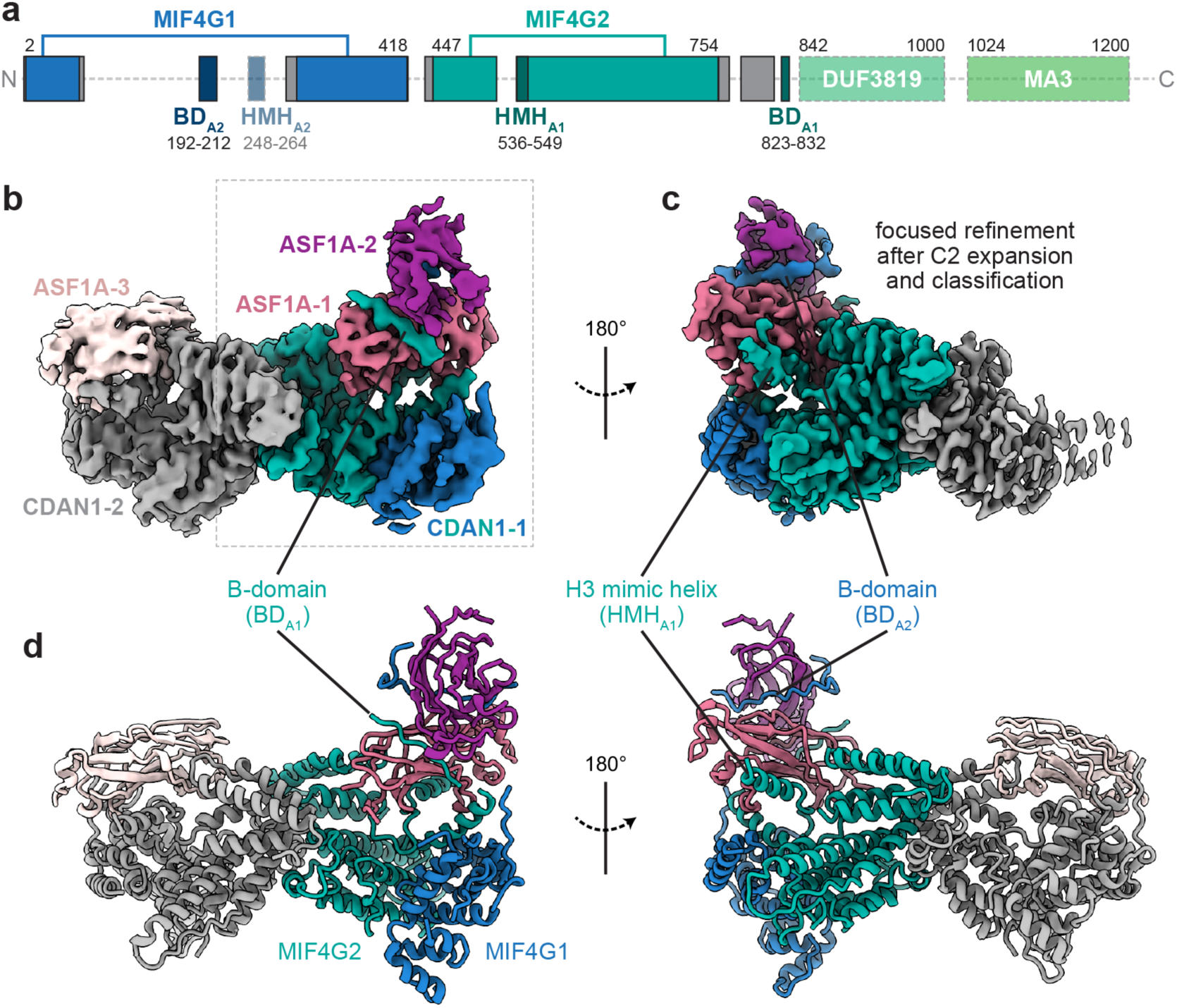
Cryo-EM structures of CDAN1-ASF1A complexes. **a,** Domain scheme of CDAN1. Gray boxes indicate structured regions modeled based on the cryo-EM maps that fall outside of defined domains. Unresolved regions are indicated by transparent boxes for regions predicted to be structured or a dashed line for unstructured regions. **b,c,** Cryo-EM maps of the C:C:A complex showing **b,** three ASF1A molecules bound to a CDAN1 dimer (contoured at 9.4σ) or **c,** focused on CDAN1 interactions with two stacked ASF1 molecules (contoured at 5.8σ) after C2 symmetry expansion and 3D classification focused on one arm of the complex. **d,** Model of a CDAN1 dimer bound to three ASF1A.

**Table 1.** Cryo-EM data collection, refinement, and validation.

|  | CDAN1 +<br>3xASF1A<br>EMD-45959<br>PDB 9CVC | CDAN1 +<br>2xASF1A (C1)<br>EMD-45960 | CDAN1 + 2xASF1A<br>(C2 expanded &<br>focused)<br>EMD-45961 |
| --- | --- | --- | --- |
| **Data collection and processing** |  |  |  |
| Magnification | 105,000 | 105,000 | 105,000 |
| Voltage (kV) | 300 | 300 | 300 |
| Electron exposure (e-/Å<sup>2</sup>) | 50.3 | 50.3 | 50.3 |
| Defocus range (μm) | 1.3 to 2.3 | 1.3 to 2.3 | 1.3 to 2.3 |
| Pixel size (Å) | 0.825 | 0.825 | 0.825 |
| Symmetry imposed | C1 | C1 | C1 |
| Initial particle images (no.) | 1,665,601 | 1,885,342 | 1,885,342 |
| Final particle images (no.) | 25,056 | 256,296 | 122,061 |
| Map resolution (Å) | 3.5 | 3.0 | 3.2 |
| FSC threshold | 0.143 | 0.143 | 0.143 |
| Map resolution range (Å) | 3.1 to 50.5 | 2.6 to 6.6 | 2.6 to 5.9 |
| **Refinement** |  |  |  |
| Initial model used | Alphafold |  |  |
| Model resolution (Å) | 4.0 |  |  |
| FSC threshold | 0.5 |  |  |
| Model resolution range (Å) | 4.0 to 45.1 |  |  |
| Map sharpening *B* factor (Å<sup>2</sup>) | -53.7 |  |  |
| Model composition |  |  |  |
| Non-hydrogen atoms | 12,529 |  |  |
| Protein residues | 1,567 |  |  |
| *B* factors (Å<sup>2</sup>) |  |  |  |
| Protein | 108.51 |  |  |
| R.m.s. deviations |  |  |  |
| Bond lengths (Å) | 0.005 |  |  |
| Bond angles (°) | 1.040 |  |  |
| Validation |  |  |  |
| MolProbity score | 2.15 |  |  |
| Clashscore | 15.17 |  |  |
| Poor rotamers (%) | 0.14 |  |  |
| Ramachandran plot |  |  |  |
| Favored (%) | 92.50 |  |  |
| Allowed (%) | 7.31 |  |  |
| Disallowed (%) | 0.20 |  |  |

Based on homology searches, CDAN1 contains two MIF4G (<u>m</u>iddle domain of eukaryotic initiation factor <u>4G</u>) domains (MIF4G1, MIF4G2), a three-helix bundle of unknown function (DUF3819), and an MA3 domain commonly found in MIF4G proteins^43,44^ (**Fig. 3a and Supplemental Fig. 5a-d**). This domain architecture, particularly the distinctive presence of the three-helix bundle, resembles the central portion of the CNOT1 protein that serves as a scaffold in the CCR4-NOT deadenylation complex^36,45–48^ (**Supplemental Fig. 5c-f).** With our cryo-EM maps, we could model both MIF4G domains of CDAN1 (**Fig. 3b-d**). This showed that MIF4G2, including a conserved central extension (residues 645-675), mediates dimerization (**Fig. 3b-d and Supplemental Fig. 5a**). MIF4G1 is split by a long loop (residues 69-284) that contains the B-domain previously identified to interact with ASF1^21^ (**Fig. 3b and Supplemental Fig. 5a**). This loop and features extending from MIF4G2 also contain additional elements that facilitate ASF1A interaction, discussed below.

Although present in the purified complex (**Fig. 2a**), CDIN1 and the C-terminal portion of CDAN1 (residues 833-1227; including the three-helix bundle and the MA3 domain that interacts with CDIN1^36,37,49^) were not visible in our cryo-EM maps. This suggests that the C-terminus of CDAN1 is flexible relative to the N-terminal core. To obtain experimental insights into this interaction, we purified the CDAN1 MA3 domain (ST-CDAN1_C_, residues 1008-1227) in complex with CDIN1 (**Supplemental Fig. 6a,b**). Single-particle cryo-EM analysis of this complex resulted in a ∼6 Å reconstruction consistent with the extensive CDAN1:CDIN1 interface predicted by Colabfold^50^ (**Supplemental Fig. 6c-e)** and recent size exclusion chromatography coupled with small angle X-ray scattering (SEC-SAXS) analysis^49^. These results support a role for CDAN1 as a scaffold that orchestrates the assembly of CDIN1 and multiple copies of ASF1 to distinct regions of the same complex.

### CDAN1 interacts with ASF1A through distinct B-domains

Surprisingly, even though CDAN1 homodimerizes, we consistently observed asymmetric assemblies during 2D and 3D classifications of our cryo-EM data. The asymmetry is explained by the association of two ASF1A molecules with one CDAN1 monomer and a single ASF1A with the other (**Fig. 3b and Supplemental Fig. 3**). This stoichiometry matches our SEC-MALS measurements indicating the predominant presence of three ASF1A in the complex (**Fig. 2c**). The two ASF1A molecules on one side of the complex appear to stack on top of each other on a platform formed primarily by MIF4G2 of CDAN1. We refer to the ASF1A proximal to the CDAN1 core as ASF1A-1 and the distal ASF1 as ASF1A-2 (**Fig. 3b-d**).

To better visualize the interactions that mediate this configuration of ASF1A-1 and ASF1A-2, we performed an intermediate 3D refinement with C2 symmetry imposed, which we then used for a symmetry expansion and 3D classification of the expanded particle set with masks focused on only one side of the dimeric complex (**Supplemental Fig. 3**). This specifically isolated classes with clear occupancy of two ASF1A that generated a 3.2 Å reconstruction after focused refinement (**Fig. 3c and Supplemental Fig. 3)**. These maps showed evidence for two distinct B-domain densities, one interacting with each ASF1A, that we refer to as BD_A1_ and BD_A2_ (**Fig. 4a-c**).

**Figure 4.**
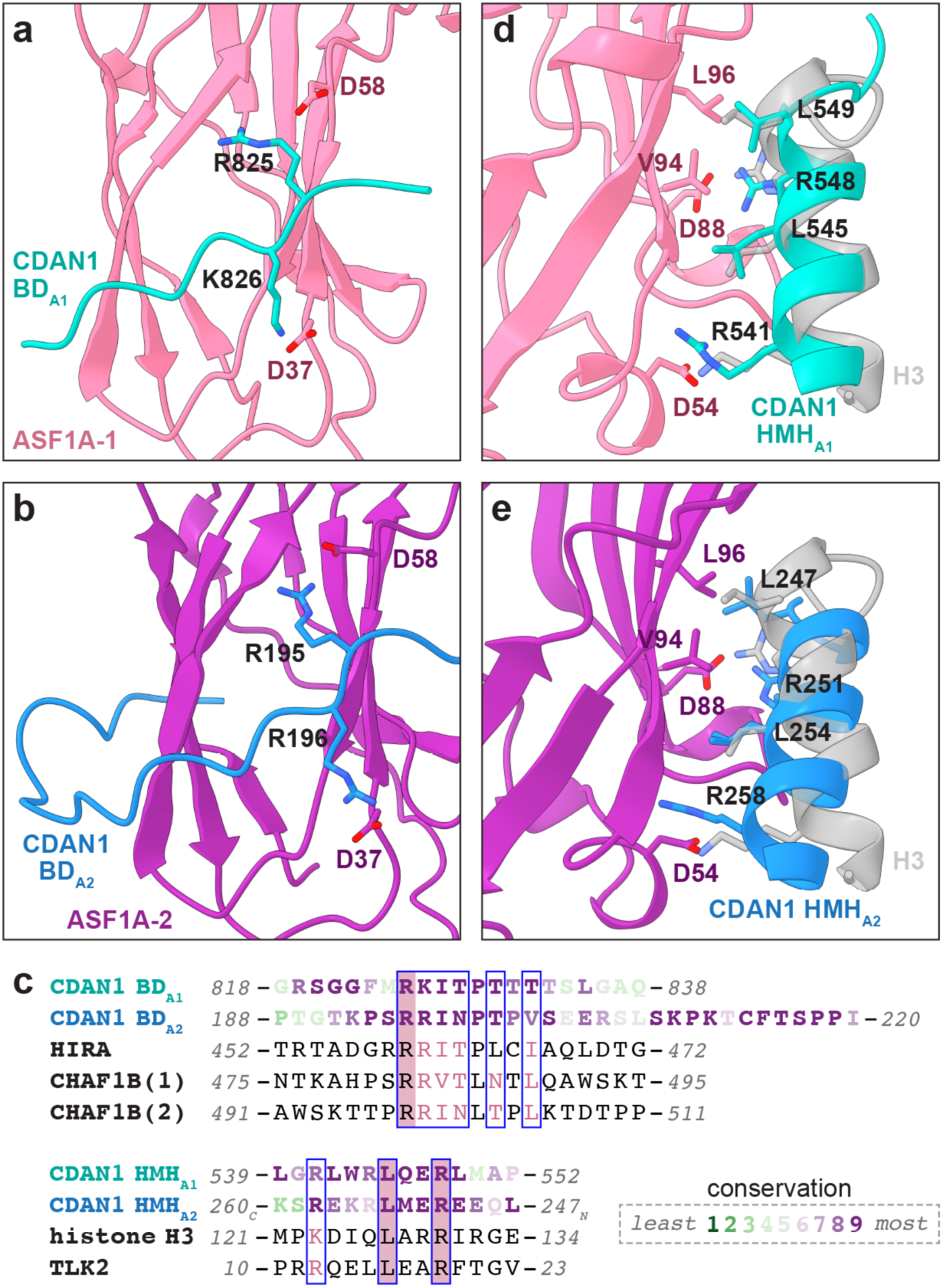
CDAN1 interacts with ASF1 through multiple motifs. **a,b,** Structural models of CDAN1 B-domain interactions with **a,** ASF1A-1 (BD_A1_) and **b,** ASF1A-2 (BD_A2_), colored as in Fig. 3. **c,** Sequence alignments of CDAN1 B-domains (BD, top) and H3 mimic helices (HMH, bottom) with other ASF1 interactors or histone H3. CDAN1 sequences are colored according to Consurf^63^ conservation values as indicated. **d,** Structural model of the CDAN1 HMH interaction with ASF1A-1 (HMH_A1_) superposed with the corresponding *X. laevis* histone H3 helix (transparent gray; residues 121-136, PDB 2IO5). **e,** Alphafold3 model of the putative CDAN1 HMH interaction with ASF1A-2 (HMH_A2_) superposed with the H3 helix (transparent gray).

Based on the organization of the structured CDAN1 domains (**Supplemental Fig. 7a**), we assigned the previously identified B-domain (residues 193-203) in the long loop extending from MIF4G2 as BD_A2_ bound to ASF1A-2 (**Fig. 4b,c**). BD_A2_ makes canonical interactions with ASF1A-2, where the consecutive basic residues R195 and R196 interact with D58 and D37 of ASF1A-2, respectively. BD_A2_ also extends into additional conserved residues that wrap around ASF1A-2, which may help stabilize the position of ASF1A-2 stacked on top of ASF1A-1.

We then searched for a sequence that can accommodate the density corresponding to BD_A1_ (**Fig. 3b**). Inspection of the CDAN1 sequence around structured regions near the BD_A1_ density identified a conserved segment (residues 823-832) consistent with a B-domain that is ideally positioned to interact with ASF1A-1 and would not be able to reach ASF1A-2 (**Fig. 3a,c and Supplemental Fig. 7a**). In this placement of BD_A1_, the consecutive basic residues R825 and K826 interact with D58 and D37 of ASF1A-1 (**Fig. 4a**) in the same way that the corresponding residues of BD_A2_ interact with ASF1A-2 (**Fig. 4b**). Hence, we have identified a second B-domain in CDAN1 that facilitates the recruitment of two ASF1A molecules.

### CDAN1 has multiple ASF1 client mimicry elements

In addition to BD _A1_, CDAN1 also interacts with ASF1A-1 through a helix (residues 536-549) that binds the same interface ASF1 uses to engage a helix of histone H3^22,51,52^ (**Fig. 4c,d**). We refer to this <u>H</u>3 <u>m</u>imic <u>h</u>elix as HMH_A1_. Like H3, HMH_A1_ of CDAN1 makes hydrophobic interactions with L96 and V94 of ASF1A-1 through L549 and L545, as well as electrostatic interactions with D54 and D88 of ASF1A-1 through R541 and R548. This client mimicry mode of ASF1 recruitment is also seen with TLK2, a kinase that phosphorylates ASF1A and ASF1B^53^.

Our assignments of HMH_A1_ and the two B-domains described above are consistent with Alphafold3 predictions of CDAN1 with two copies of ASF1A, including the observation that BD _A1_ and HMH_A1_ bind to the same ASF1A molecule (**Supplemental Fig. 7b**). Notably, Alphafold3 also predicts that another CDAN1 helix (residues 248-264) binds the H3-binding interface of the other ASF1A molecule associated with BD_A2_ (**Supplemental Fig. 7b**). The placement of this helix is consistent with density at this interface of ASF1A-2 visible in unsharpened cryo-EM maps of the C:C:A complex (**Supplemental Fig. 7c**). We therefore assign this helix as HMH_A2_ (**Fig. 3a** and **4c,e**). Compared to H3 and HMH_A1_, the predicted N-to-C orientation of HMH_A2_ is reversed but still positions conserved amino acid to make key interactions with the ASF1A interface (**Fig. 4c,e**). In this position, L247 and L254 of HMH_A2_ would satisfy hydrophobic interactions with L96 and V94 of ASF1A-2, and R251 and R258 of HMH_A2_ would interact with D88 and D54 of ASF1A-2.

CDAN1 interactions with ASF1A-1 and ASF1A-2 would compete not only with H3 binding but also potentially with an interaction between ASF1A and the C-terminal tail of histone H4^52^ (**Supplemental Fig. 7d**). Specifically, a loop (residues 469-476) associated with CDAN1 MIF4G2 would clash with an H4 tail positioned on ASF1A-1, and the extended BD_A2_ loop would clash with an H4 tail positioned on ASF1A-2 (**Supplemental Fig. 7d**). Altogether, our structural analyses show that a single CDAN1 molecule can recruit two ASF1A molecules through two distinct B-domains and two H3 mimic helices that would interfere with the histone chaperoning activity of both ASF1A proteins.

### ASF1A and ASF1B have different CDAN1 binding requirements

ASF1A and ASF1B are over 70% identical (**Supplemental Fig. 8a**). The most divergent portions of the two paralogs are their N-terminal domains (NTD, residues 1-27) and unstructured C-terminal tails that are not resolved in our cryo-EM structures (**Supplemental Fig. 8a,b**). Interestingly, mutation of key residues in the previously identified BD_A2_ sequence or the well-resolved HMH_A1_ revealed that ASF1B was more dependent than ASF1A on HMH_A1_ for interaction with ST-CDAN1 (**Fig. 5a,b**). While wildtype (WT) CDAN1 pulled down both ASF1A and ASF1B as expected (**Fig. 5b**, lane 7), H* CDAN1 carrying mutations in HMH_A1_ copurified ASF1A but not ASF1B (**Fig. 5b**, lane 8). ASF1A and ASF1B could both bind B* CDAN1 carrying mutations in BD_A2_ (**Fig. 5b**, lane 9), while mutating both HMH_A1_ and BD_A2_ abolished binding to both ASF1 paralogs (**Fig. 5b**, lane 10). In all cases, CDIN1 association with the CDAN1 variants was not impacted, indicating that these mutations do not generally disrupt CDAN1 structure. Thus, ASF1B but not ASF1A is nearly exclusively reliant on the proximal HMH for stable binding to CDAN1.

**Figure 5.**
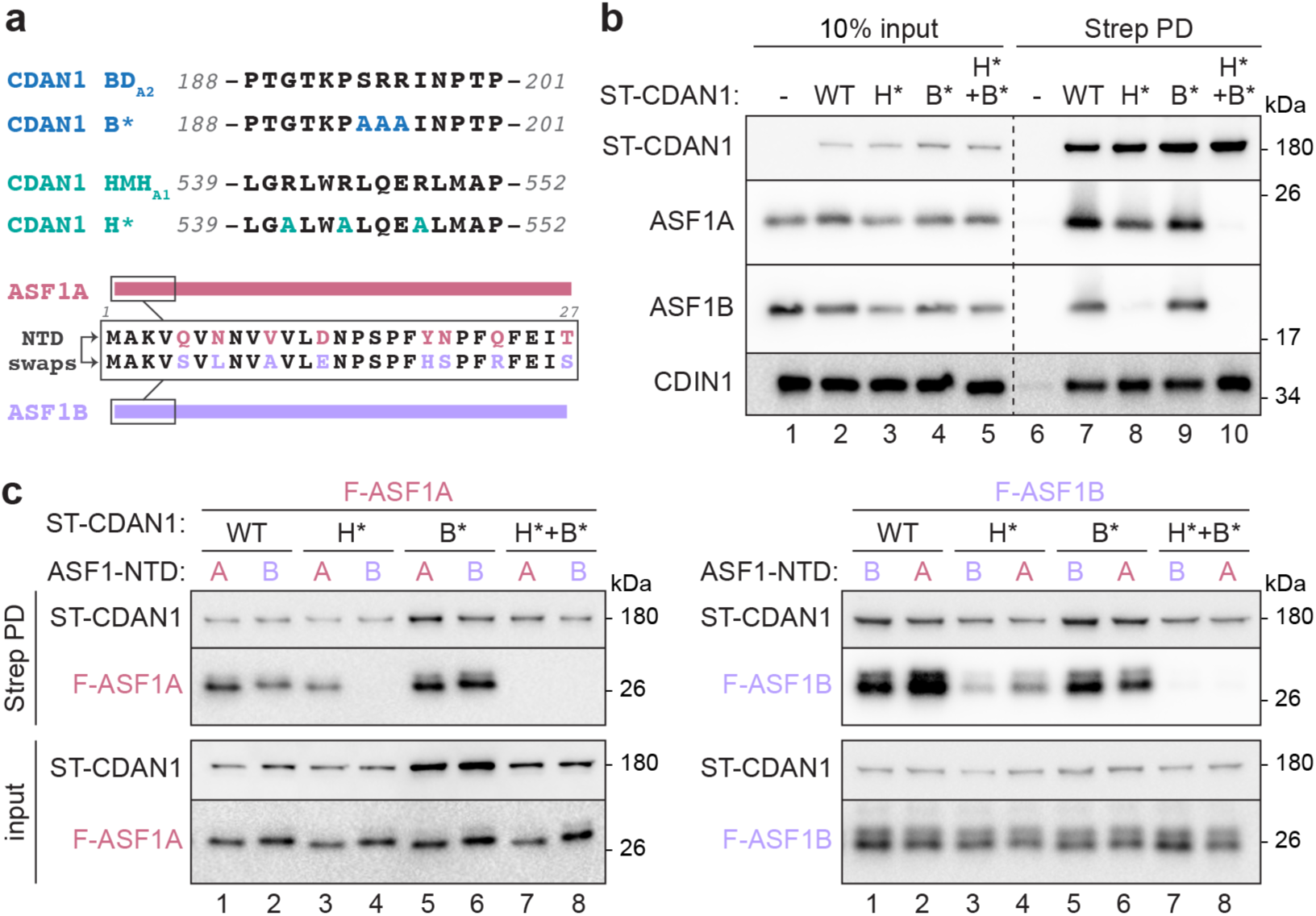
ASF1A and ASF1B have different CDAN1 binding requirements. **a,** Mutations of key residues in the canonical CDAN1 B-domain (BD_A2_ mutated to B*) and H3 mimic helix (HMH_A1_ mutated to H*) (top) and scheme of ASF1A and ASF1B N-terminal domain (NTD) swaps analyzed. **b,** ASF1B interaction with CDAN1 is more reliant on HMH_A1_ than ASF1A. Flp-In 293 T-REx cells transiently transfected with Strep-tagged CDAN1 (ST-CDAN1) variants without or with mutations in BD_A2_ (B*) and/or HMH_A1_ (H*) were lysed (input) and subjected to Strep-Tactin pulldowns (PD) before analysis by SDS-PAGE and immunoblotting; representative of 3 independent replicates. **c,** The indicated ST-CDAN1 variants were co-transfected with FLAG-tagged ASF1A (F-ASF1A, left) or ASF1B (F-ASF1B, right) with the indicated N-terminal domain (NTD). Cell lysates before (input) or after Strep PD were analyzed by SDS-PAGE and immunoblotting; representative of 2 independent replicates.

The histone chaperone HIRA has also been reported to bind ASF1A with higher affinity, in part by distinguishing the variable NTD of ASF1A and ASF1B that lie proximal to the B-domain interface^28^ (**Supplemental Fig. 8b**). To investigate if the NTD of ASF1A and ASF1B also contribute to their different binding preferences to CDAN1, we generated NTD swaps of each ASF1 paralog and analyzed their association with ST-CDAN1 variants by co-transfection and Strep-Tactin pulldowns (**Fig. 5a,c**). This revealed that swapping the NTD of ASF1A to that of ASF1B diminishes binding to WT CDAN1 (**Fig. 5c**, left, compare lanes 1 and 2) and abrogates binding to H* CDAN1 (**Fig. 5c**, left, compare lanes 3 and 4). Reciprocally, replacing the NTD of ASF1B with that of ASF1A enhanced binding to both WT and H* CDAN1 (**Fig. 5c**, right, lanes 1-4). These results indicate that if the HMH_A1_ interface is disrupted, a compensatory interaction can occur between CDAN1 and ASF1A but not ASF1B. This paralog-specific interaction appears to at least partially involve BD_A2_ of CDAN1 and the NTD of ASF1A but not of ASF1B.

Consistent with this interpretation, H*+B* CDAN1 harboring mutations in both HMH_A1_ and BD _A2_ failed to bind any ASF1 variant (**Fig. 5c**, lanes 7-8). In addition, B* CDAN1 pulled down similar amounts of ASF1A without or with an NTD swap (**Fig. 5c**, left, lanes 5 and 6), supporting the interpretation that the B-domain contributes to the differential binding behaviors associated with the two NTDs. Considered together, our findings support a model in which HMH_A1_, which packs against the structured MIF4G2 domain, forms a major docking site on CDAN1 for free ASF1 that is not bound to H3-H4. When bound to HMH _A1_, ASF1 can no longer bind H3-H4 and is further secured by BD_A1_. Independently, the long CDAN1 loop emerging from MIF4G1, which contains both BD_A2_ and the putative HMH_A2_, is able to capture another ASF1 molecule via interactions that may be favored for ASF1A over ASF1B and that ultimately establish the stacked conformation of ASF1A observed in our structures.

## Discussion

Our findings reveal how CDAN1 is able to sequester and inhibit the chaperone function of multiple ASF1 molecules. A recent study^51^ also reported CDAN1 binding to ASF1A via a H3 mimic helix (i.e. HMH_A1_) and a B domain (i.e. BD_A2_). In addition, we identify a previously unappreciated second B-domain (BD_A1_) on CDAN1 and another putative H3 mimic helix (HMH_A2_) that together permit one copy of CDAN1 to inhibit the chaperoning interface of two ASF1A. Although a CDAN1 dimer has four putative ASF1 binding sites, we do not observe binding of a fourth ASF1 molecule. This may be due to conformational restraints that limit the stable ‘stacking’ of the distal ASF1A-2 on both arms of the dimer. Alternatively, regions of CDAN1 that are not resolved in our cryo-EM maps, such as the three-helix bundle or MA3 domain, may sterically hinder recruitment of a fourth ASF1.

The simultaneous use of multiple B-domains and H3 mimic helices by CDAN1 to suppress ASF1 histone chaperoning activity is unique among ASF1 interactors. ASF1 can interact with other chaperones, such as HIRA and the p60 subunit of CAF-1 (CHAF1B), via a B-domain^28,29,31,54^. However, unlike CDAN1, these factors engage ASF1 bound to an H3-H4 heterodimer^55^ primarily in the nucleus as part of productive nucleosome assembly pathways during replication-independent (HIRA) or replication-dependent (CAF-1) deposition^56,57^. Other factors have also been reported to bind ASF1 through the H3-binding interface. Similar to CDAN1, yeast Rad53 and Rtt109 use both a B-domain and an additional region to bind ASF1 and compete with histone binding^30,32^. However, these interactions appear to occur only in yeast^30,58^. TLK2 also uses a H3 mimic helix to engage ASF1 and to mediate ASF1 phosphorylation, but this is suggested to promote histone binding and nucleosome assembly^53^. CDAN1 is therefore distinctive in its ability to simultaneously engage multiple ASF1 molecules and divert them from promoting histone deposition.

Although CDAN1 binds both ASF1A and ASF1B, there are different requirements for the interaction of each paralog. In comparison, CAF-1 binds both ASF1 paralogs^59^, while HIRA specifically binds ASF1A^28^. CDAN1 may therefore impact the binding of ASF1 chaperones to specific paralogs to different extents. In most dividing cells, the expression of CDAN1 is low relative to other interactors. For example, in HEK293T cells, CAF-1 is approximately equimolar with ASF1A or ASF1B, which are each ∼10-fold more abundant than CDAN1^60^. Similarly, HIRA is several-fold more abundant than CDAN1. Thus, although CDAN1 can bind multiple ASF1 molecules, it is unlikely to engage a significant proportion of ASF1 in most cells, even though the essential nature of CDAN1 suggests it nonetheless has a critical function in most cell types.

However, if the balance of the levels of ASF1 and its interactors shifts, CDAN1 could potentially assume an outsized role by quantitatively competing with histone deposition pathways. Such dynamics may arise during erythropoiesis, when maturing erythroblasts undergo extensive proteome remodeling and chromatin condensation in preparation for organellar clearance^61^. At the proerythroblast stage, the levels of ASF1 interactors are altered, with approximately equal levels of CAF-1 and CDAN1 that are both expressed in ten-fold excess of HIRA^62^. Under such conditions, CDAN1 may be capable of inhibiting a substantial proportion of ASF1 that is significant enough to cause pathogenic defects in chromatin condensation when its function or expression is disrupted through mutations. The molecular level insights uncovered in our study will drive forward our understanding of these functions of CDAN1 by facilitating detailed dissections of this complex in different physiological contexts.

## Acknowledgements

Cryo-EM screening and data collection were performed at the Harvard Center for Cryo-Electron Microscopy (HC2EM). Data processing was supported by SBGrid. Light microscopy was performed at the Core for Imaging Technology and Education (CITE) at Harvard Medical School (HMS). SEC-MALS was performed at the Center for Macromolecular Interactions (CMI) at HMS. We thank H. Chino for guidance with endogenous tagging and help with cell sorting, M. Yip for guidance with cryo-EM sample preparation and processing, M. McKenna and D. Sherpa for critical reading, and Shao lab members for useful discussions. This work was supported by NIH F31HL157976 (SFS), NIH DP2GM137415 (SS), and a Packard Fellowship (SS).

## Competing Interests

The authors declare no competing interests.

## Author Contributions

SFS conceived the project and performed all investigations. SS supervised the project and acquired funding. SFS and SS wrote the paper.

## Supplemental Figures

**Supplemental Figure 1.**
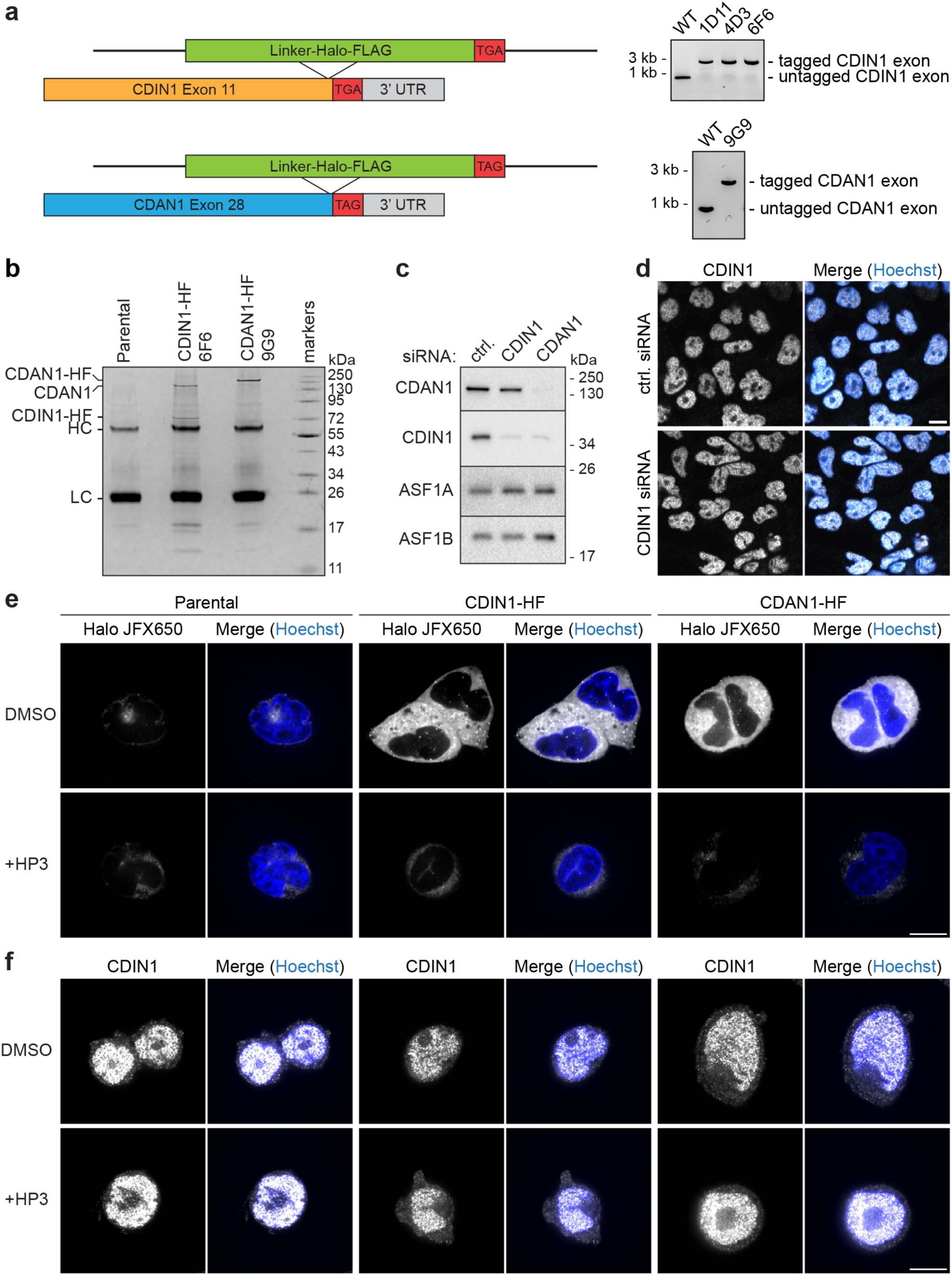
Characterization of endogenous CDAN1 and CDIN1. **a,** Scheme (left) for endogenously tagging CDIN1 and CDAN1 with a C-terminal HaloTag-FLAG (HF) tag and PCR validation of knock-in clones (right). **b,** SDS-PAGE and Coomassie staining of anti-FLAG resins after immunoprecipitations of lysates from the indicated parental, CDIN1-HF, and CDAN1-HF Flp-In 293 T-REx cells. HC – heavy antibody chain, LC – light antibody chain. Note: CDIN1-HF copurifies an approximately equimolar amount of endogenous CDAN1; endogenous CDIN1 between LC and HC. **c,** SDS-PAGE and immunoblotting of lysates of Flp-In 293 T-REx cells treated with control (ctrl.) siRNAs or siRNAs against CDIN1 or CDAN1. Note: knocking down CDAN1 destabilizes CDIN1, but not vice versa; representative of 3 independent replicates. **d,** Immunofluorescence using anti-CDIN1 antibody of Flp-In 293 T-REx cells treated with ctrl. siRNA (siNeg) or siRNA against CDIN1, co-stained with Hoechst. Note: the nuclear signal observed with the anti-CDIN1 antibody does not decrease after knocking down CDIN1. **e,** Live-cell imaging of the indicated cells labeled with the JFX650 HaloTag ligand after treatment with DMSO or 500 nM HaloPROTAC3 (HP3) for 48 hr. Note: the specific cytosolic HaloTag signal in CDIN1-HF and CDAN1-HF cells is abolished after HP3 treatment. **f,** Immunofluorescence of cells from **e** after fixation using anti-CDIN1 antibody. Note: signal from the antibody is observed in all conditions and does not decrease in any cell line after HP3 treatment. Scale bar, 10 µm for all images.

**Supplemental Figure 2.**
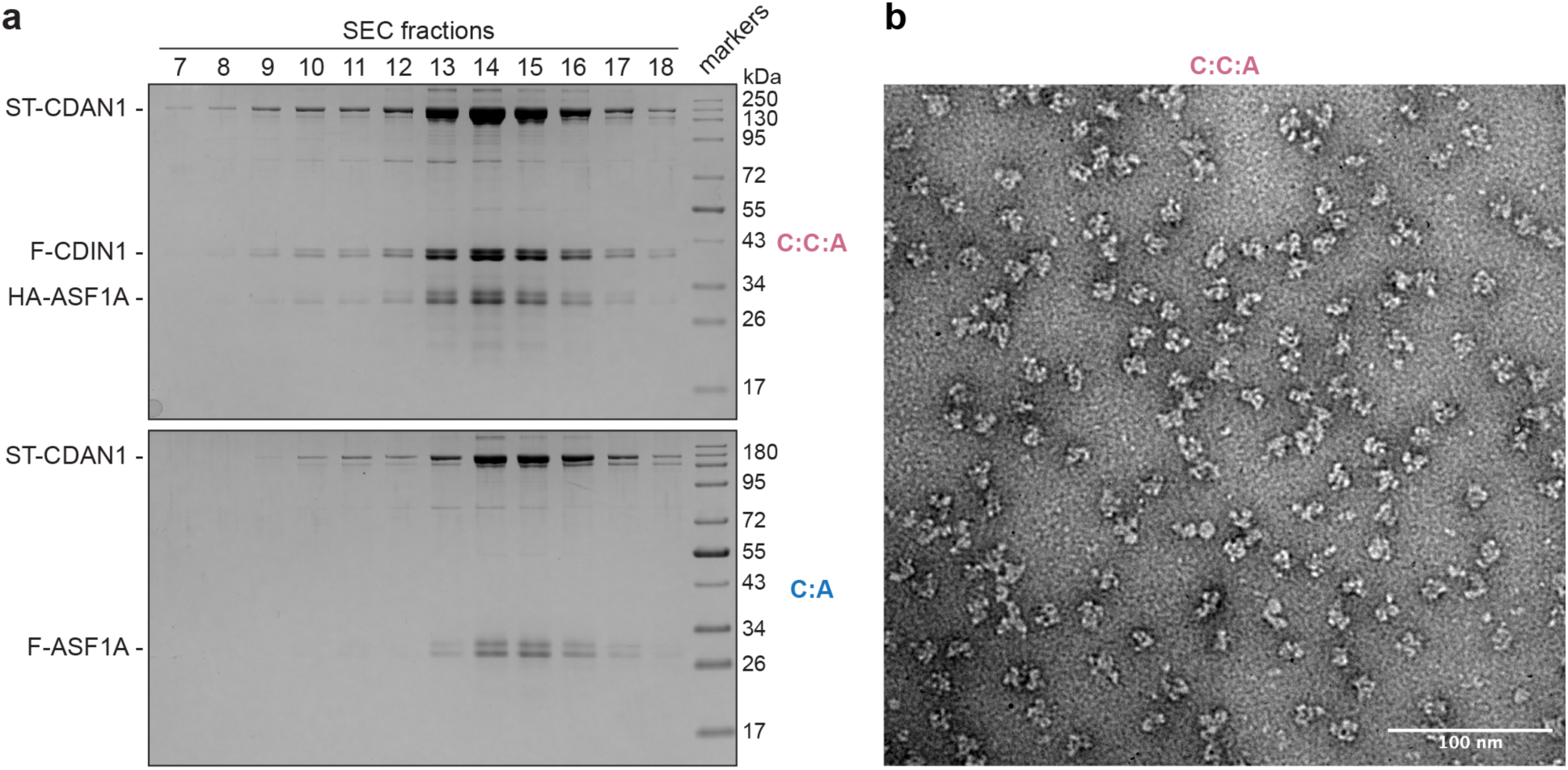
Characterization of purified CDAN1 complexes. **a,** Size exclusion chromatography fractions of purified CDAN1-CDIN1-ASF1A (C:C:A; top) or CDAN1-ASF1A (C:A; bottom) analyzed by SDS-PAGE and Coomassie staining show comigration of the complex components. **b,** Representative negative stain EM image of the C:C:A complex.

**Supplemental Figure 3.**
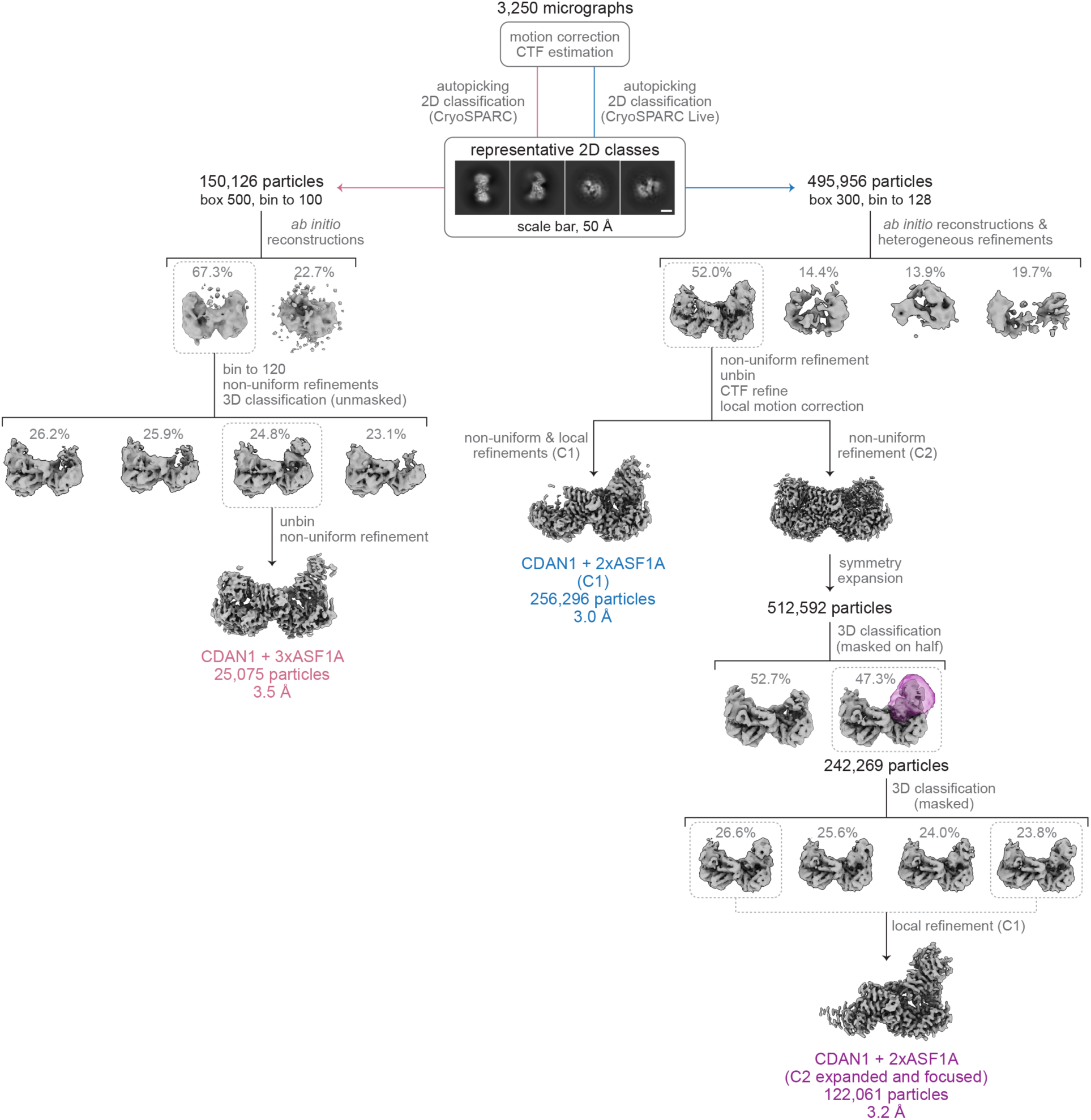
Cryo-EM data processing summary. Overview of single-particle cryo-EM data processing and classification pipeline. The map with three ASF1A molecules was obtained from a separate autopicking and 2D classification pipeline using a larger box size (left) than the other maps, which were obtained from initial processing in CryoSPARC Live. Because asymmetry was consistently observed in the maps, the refinement with C2 symmetry was only used for symmetry expansion. The expanded particle set was then subjected to 3D classification masked on elements on one half of the map to obtain higher resolution insights into how two ASF1A molecules are stacked on one side of the CDAN1 dimer.

**Supplemental Figure 4.**
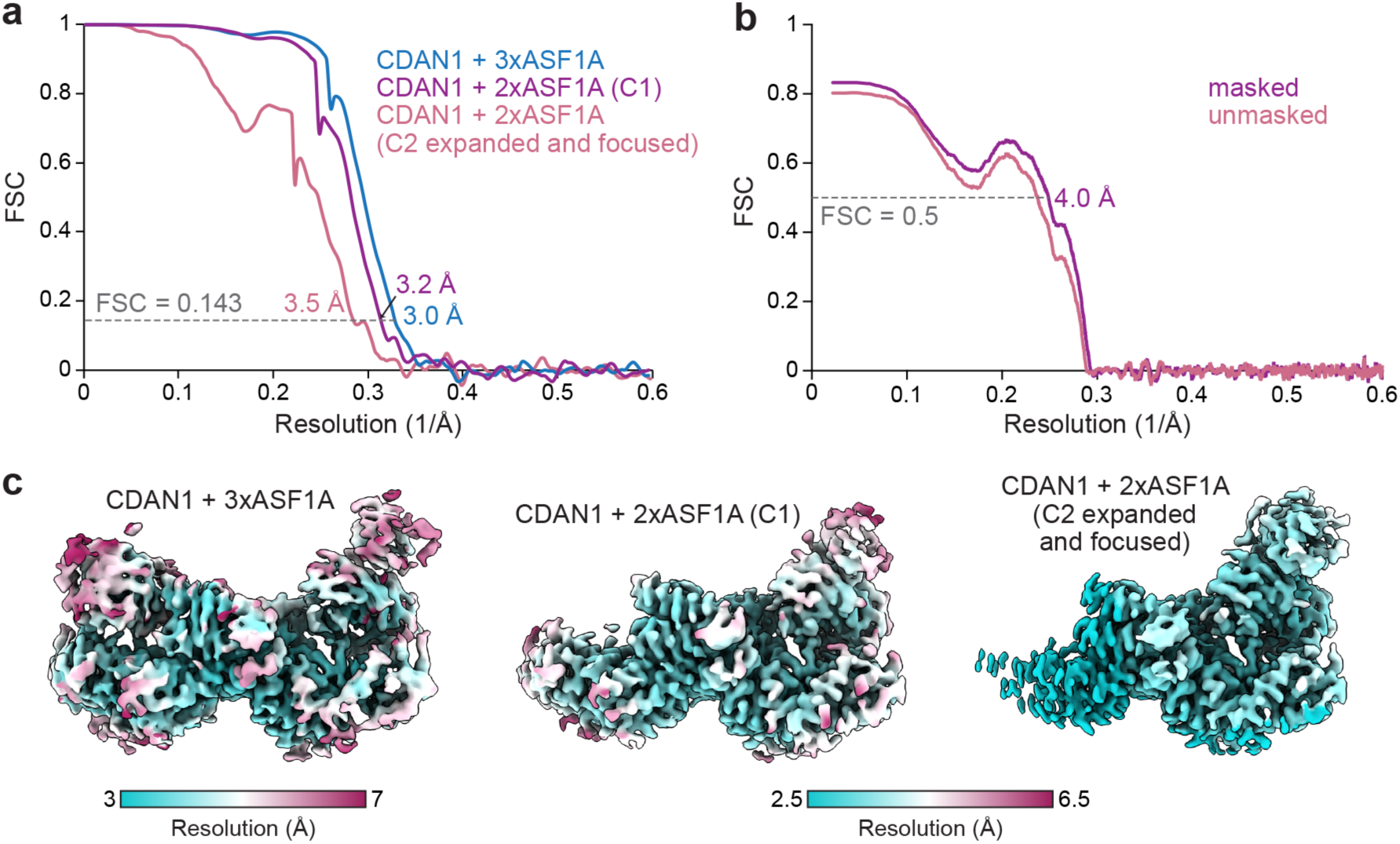
Quality of maps and models. **a,** Fourier shell correlation (FSC) vs. resolution (1/Å) curves for the indicated cryo-EM maps. **b,** Model vs. map FSC curves. **c,** The indicated cryo-EM maps colored by local resolution.

**Supplemental Figure 5.**
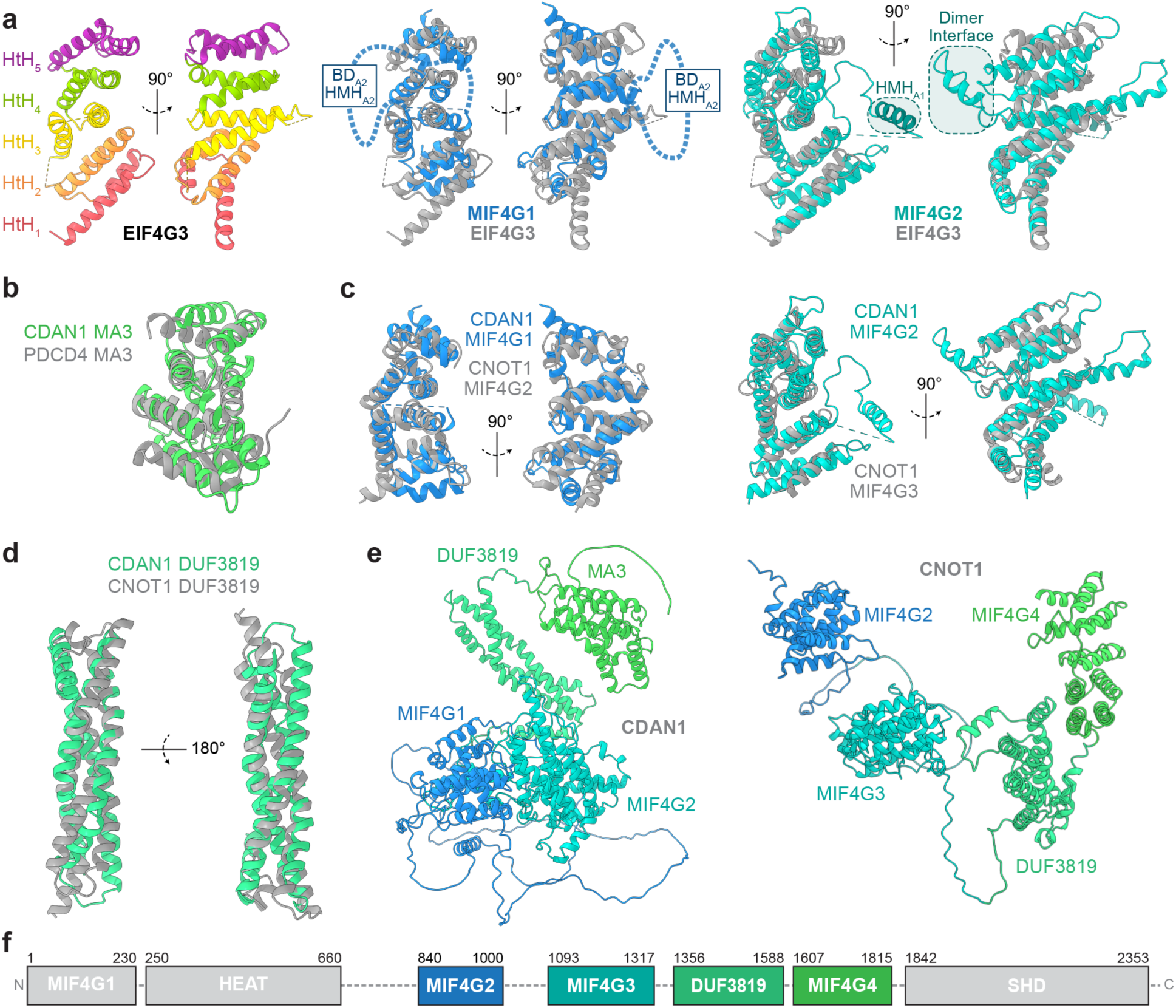
CDAN1 domain analysis. **a,** Canonical MIF4G domain of eIF4G3 (PDB 1HU3) with each helix-turn-helix (HtH) motif colored separately (left) or superposed (gray) with CDAN1 MIF4G1 (middle; blue) or CDAN1 MIF4G2 (right; teal). **b,** Canonical MA3 domain of PDCD4 (PDB 2RG8, gray) superposed with the CDAN1 MA3 domain (green; Dali Z-score 7.8). **c,** Superposition of CDAN1 MIF4G1 (blue) with CNOT1 MIF4G2 (gray, left) or of CDAN1 MIF4G2 (teal) with CNOT1 MIF4G3 (gray, right). **d,** Superposition of the three-coil bundle domain of unknown function (DUF3819) predicted to be present in both CDAN1 (sea green) and CNOT1 (gray). **e,** Alphafold2 model of CDAN1 (left) or the central region of CNOT1 (right) colored by domain. **f,** Domain scheme of CNOT1 with the central region similar to CDAN1 colored as in **e**.

**Supplemental Figure 6.**
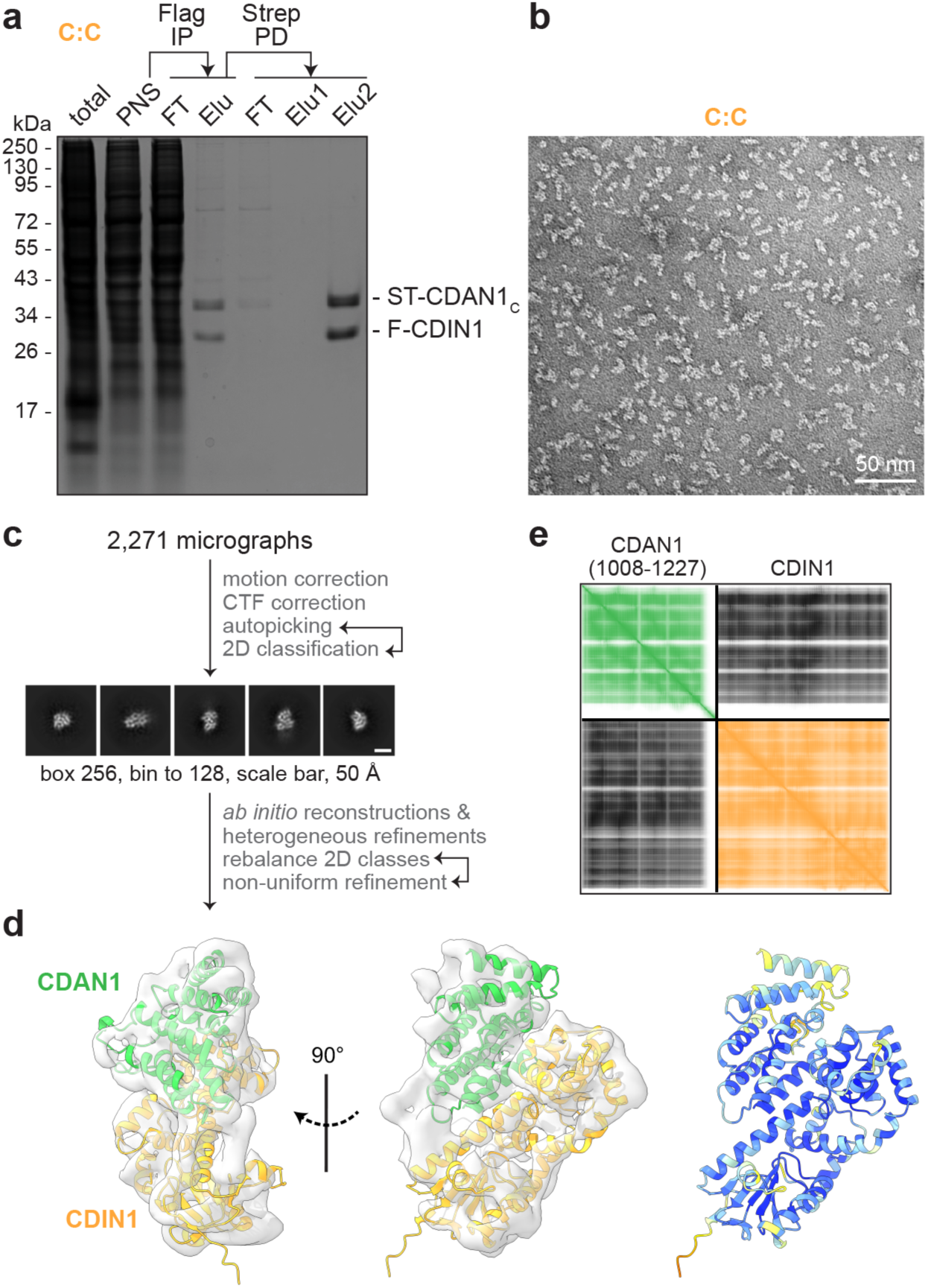
Analysis of the CDAN1-CDIN1 interaction interface. **a,** The Strep-tagged MA3 domain of CDAN1 (ST-CDAN1_C_;, residues 1008-1227) was co-expressed with FLAG-tagged CDIN1 (F-CDIN1) in Expi293 cells by transient transfection. The cells were then lysed, and the post-nuclear supernatants (PNS) were subjected to anti-FLAG immunoprecipitation (FLAG IP) followed by Strep-Tactin pulldowns (Strep PD) to purify the CDAN1_C_-CDIN1 (C:C) complex. The total lysate, PNS, flow-through (FT), and elution (Elu) samples were analyzed by SDS-PAGE and Coomassie staining. **b,** Representative negative stain EM image of the C:C complex. **c,** Summary of cryo-EM data processing scheme for the C:C complex. **d,** Colabfold model of the C:C complex colored by chain (CDAN1 – green, CDIN1 – light orange) or pLDDT values docked into a ∼6 Å cryo-EM map (left, transparent gray)**. e,** Predicted alignment error (PAE) plot of the Colabfold^50^ prediction of the C:C complex.

**Supplemental Figure 7.**
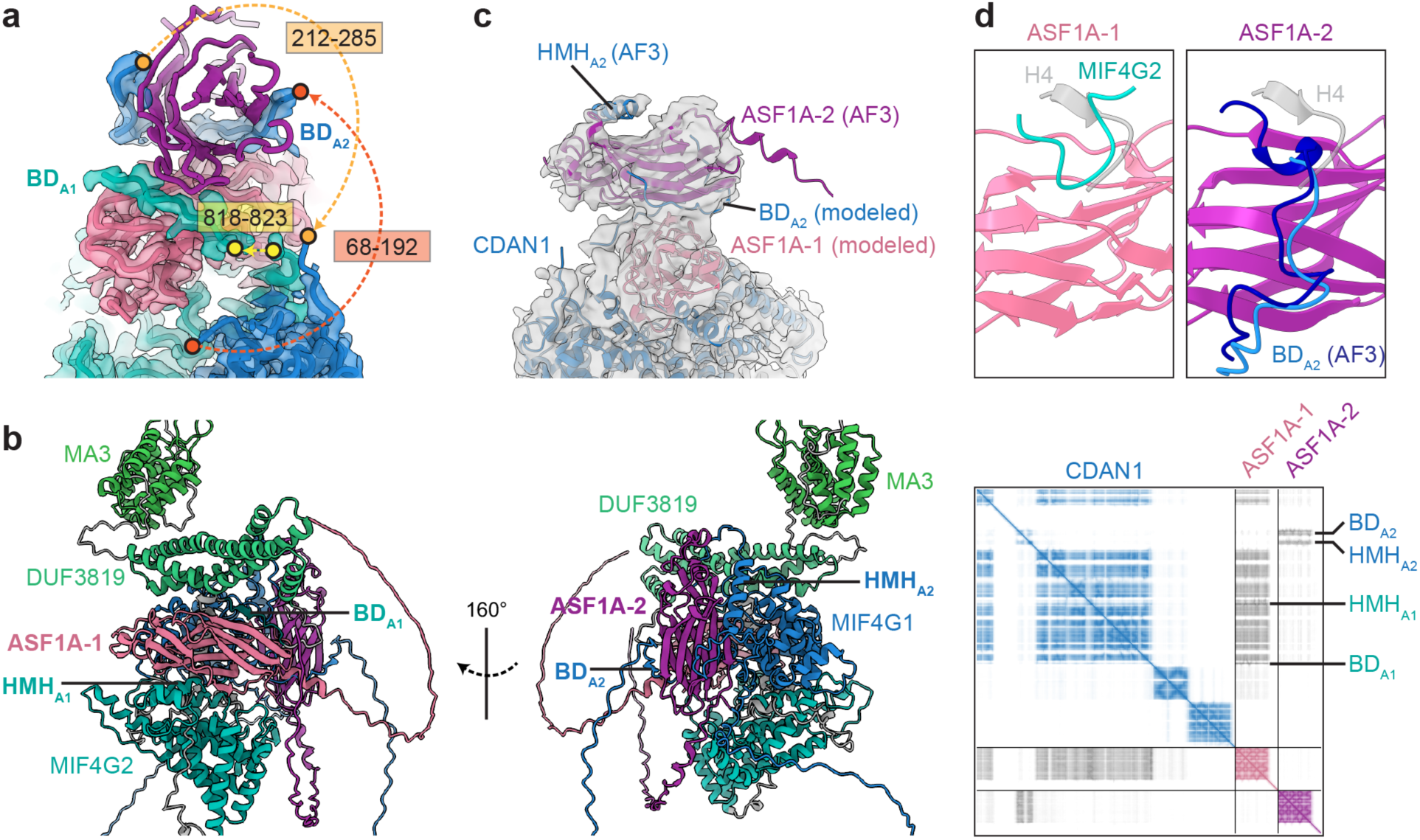
CDAN1 interactions with ASF1A. **a,** CDAN1 B-domain assignments based on connectivity. Overlay of the structural model of CDAN1 bound to two stacked ASF1A molecules with the focused cryo-EM map contoured at 4.4σ; density for ASF1A-2 is omitted for clarity. A >200 amino acid loop (residues 69-284) extending from CDAN1 MIF4G2 (blue) harboring the previously identified B-domain sequence (BD_A2_; residues 192-212 are modeled) and a putative H3 mimic helix (see **c**) is sufficiently long to engage the distal ASF1A-2 (purple). Dark orange and light orange dotted lines indicate how BD_A2_ links to the structured portion of MIF4G2. In contrast, the newly identified BD_A1_ sequence (residues 823-832) can only reach to ASF1A-1 (pink) from the modeled structured domain following MIF4G (yellow dotted line). **b,** Alphafold3 (AF3) model (left) and of CDAN1 with two ASF1A molecules, colored as in Fig. 3, and associated predicted alignment error (PAE) plot (right) predict distinct B-domain (BD) and H3 mimic helix (HMH) interactions with each ASF1A. **c,** The AF3 model of ASF1A bound to the putative HMH_A2_ spanning residues 247-260 of CDAN1 was superposed to ASF1A-2 of our structural model as in Fig. 3d and docked into the unsharpened CDAN1 + 3xASF1A map contoured at 6.8σ. Note: unmodeled density corresponding to the position of HMH_A2_ predicted by AF3. **d,** Structural models of ASF1A-1 and a CDAN1 MIF4G2 loop (residues 469-476; left) or the AF3 model of ASF1A-2 engaged with the extended BD_A2_ modeled in our cryo-EM structure (residues 201-212; light blue) or residues 201-218 (dark blue) in the AF3 model (right), both superposed with PDB 2IO5 showing the position of the C-terminal tail of histone H4 (residues 96-101; transparent gray). Note: clash of CDAN1 elements with the H4 tail position on both ASF1A-1 and ASF1A-2.

**Supplemental Figure 8.**
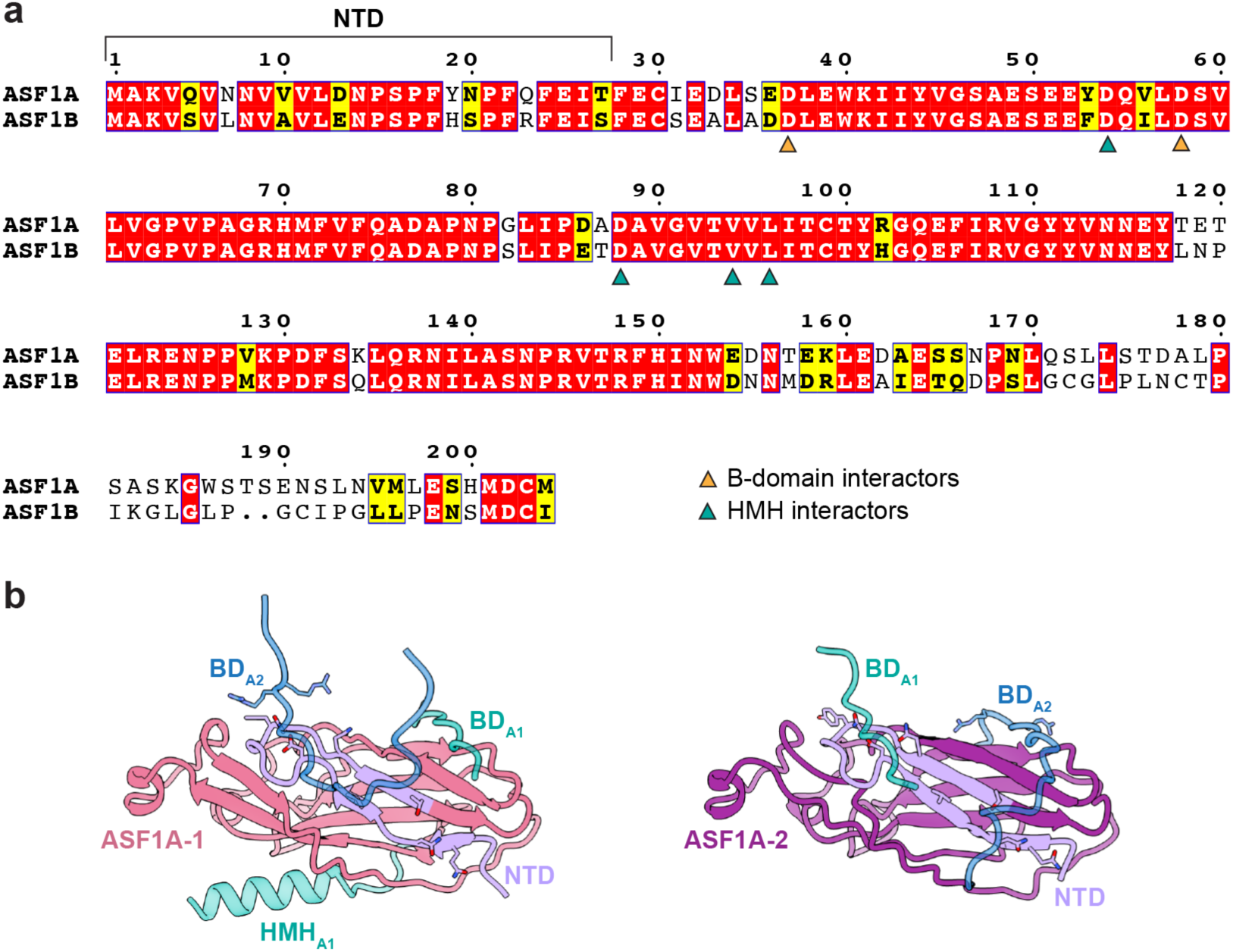
ASF1A and ASF1B comparisons. **a,** Sequence alignment of ASF1A and ASF1B. Residues that mediate interaction with B-domains (orange triangle) and with histone H3 or H3 mimic helices (HMH; teal triangles). **b,** Placement of CDAN1 B-domains (BD) and the proximal HMH relative to ASF1A-1 (left) and ASF1A-2 (right). The N-terminal domain (NTD; lavender) on each ASF1A is indicated.

## Materials and Methods

### Plasmids and antibodies

The initial construct encoding HaloTag was a gift from the Adelman lab^64^. The CDAN1 cDNA was purchased from Horizon Discovery/Dharmacon (MHS6278-202833430); a single point mutation (S421F) was corrected by Gibson assembly. cDNA sequences for ASF1A, ASF1B, and CDIN1 were synthesized by IDT. cDNA sequences were cloned into a pCDNA3.1 mammalian expression vector encoding an N-terminal 3xFLAG, 3xHA, or 3xStrep-II-TEV tag, where indicated. Mutagenesis was performed with either Gibson assembly or Phusion mutagenesis to generate <u>Strep-CDAN1 H*</u> (R541A/R544A/R548A), <u>Strep-CDAN1 B*</u> (S194A/R195A/R196A), <u>Strep-CDAN1 H*+B*</u> (S194A/R195A/R196 + R541A/R544A/R548A),

FLAG-ASF1A>N (residues 1-27 from ASF1B swapped with residues 1-27 of ASF1A), FLAG-ASF1B>N (residues 1-27 from ASF1A swapped with residues 1-27 of ASF1B), and Strep-CDAN1_C_ (residues 1008-1227).

For endogenous tagging, donor templates were constructed in a pKI plasmid (a gift from H. Chino) encoding 500 bp homology arms flanking the HaloTag-FLAG insert followed by a stop codon. The PAM site was mutated in the template to prevent self-cleavage. Homology arm sequences were amplified from gDNA for both CDAN1 and CDIN1. Guide RNAs targeting CDAN1 (CTGGTGCAATGCCCAAGGCA) or CDIN1 (GCATAAGGTGACAATGTTCG) were designed using Benchling (https://benchling.com) and inserted into the pX459 plasmid^65^ using restriction enzyme cloning.

Antibodies used for blotting: HRP-conjugated anti-FLAG M2 (Sigma, A8592, 1:10,000), HRP-conjugated Strep-Tactin (Bio-rad, 1610381, 1:5,000), HRP-conjugated anti-HA (Cell Signaling Technology, 2999S, 1:5,000), anti-CDIN1 (Abcam, ab215190, 1:1,000), anti-CDAN1 (Bethyl, A304-952A, 1:1000), anti-histone H3 (Proteintech, 17168-1-AP, 1:5000), anti-histone H4 (Proteintech, 16047-1-AP, 1:1000), anti-DNAJC9 (Proteintech, 25444-1-AP, 1:1000), anti-ASF1A (Cell Signaling Technology, 2990S, 1:1000), and anti-ASF1B (Proteintech, 22258-1-AP, 1:1000). Antibodies used for immunofluorescence: CDIN1 (Atlas Antibodies, HPA061023, 1:50).

### Cell culture and cell line generation

Flp-In 293 T-REx cells were cultured in Dulbecco’s Modified Eagle’s Medium (DMEM) supplemented with 10% fetal bovine serum (FBS) at 37°C and 5% CO_2_. Expi293F cells were cultured in Expi293 medium (Gibco A1435101) at 37°C and 8% CO_2_ with shaking at 120 r.p.m.

To generate endogenously tagged cell lines, Flp-In 293 T-REx cells were reverse co-transfected with pKI and individual pX459 plasmids using TransIT-293 (Mirus) according to the manufacturer’s instructions in media containing 690 nM Alt-R™ HDR Enhancer V2 (IDT, 10007910). After 48 hr, cells were placed under 2 μg/mL puromycin (Gibco, A11138) selection for an additional 48 hr. After selection, cells were labeled with 100 nM HaloTag TMR ligand (Promega, G8252) for 30 min at 37°C to detect insertion of the donor template. Single clones with positive fluorescent signal were sorted into 96-well plates using a Sony SH800 Sorter. Cell lysates were isolated from each colony and tagging was confirmed by immunoblotting to detect the anticipated size shift of the endogenous protein. The epitope recognized by the CDAN1 antibody is blocked by the HaloTag. All clones were also validated by PCR and Sanger sequencing of the endogenous locus.

For siRNA treatment, Flp-In 293 T-REx cells were reverse transfected with 10 nM siRNA (siNeg, siCDAN1, or siCDIN1 [Horizon Discovery/Dharmacon, D-001810-10-20, L-015478-00-0005, or L-014906-02-0005]) using Lipofectamine RNAiMAX (Invitrogen, 13778-150) according to the manufacturer’s instructions. For Supplemental Fig. 1d, cells were plated in a 12-well glass bottom plate (Mattek, NC0190134). After 48 hr, cells were either lysed for immunoblotting or washed with phosphate-buffered saline (PBS) and fixed for immunofluorescence. For targeted degradation, cells were treated with either 500 nM HaloProtac3 (HP3; a gift from the Adelman lab or purchased from Promega, GA3110) or DMSO/Ent-HP3 (Promega, GA4110) for the specified amount of time.

### Co-immunoprecipitations

In general, Flp-In T-REx 293 cells were co-transfected with 1 µg plasmid per construct in a 6-well plate using TransIT-293 according to the manufacturer’s instructions. 16–24 hr after transfection, the cells were collected in cold PBS, lysed in lysis buffer (50 mM HEPES pH 7.5, 100 mM KOAc, 2.5 Mg(OAc)_2_, 1% digitonin with 1 mM dithiothreitol [DTT], 1× cOmplete protease inhibitor cocktail [PIC; Roche, 1187358] with or without 1× PhosStop (Roche)) for 10 min on ice, and clarified by centrifugation at 21,130×*g* for 10 min at 4°C. Lysis buffer for Strep-Tactin pulldowns (PD) also contained BioLock (IBA, 2–0205). Normalized lysates were added to immunoprecipitation (IP) buffer (50 mM HEPES pH 7.5, 100 mM KOAc, 2.5 Mg(OAc)_2_, 1% Triton X-100) containing either anti-FLAG M2 agarose (Sigma, A2220) or Strep-Tactin Sepharose HP resin (Cytiva, 28-9355-99) and incubated while rotating for 1 hr at 4°C. After rotation, samples were washed 3 times with IP buffer and eluted in protein sample buffer.

To immunoprecipitate endogenously tagged CDAN1 or CDIN1, 12 15 cm dishes of each cell line were lysed in 12 mL IP buffer supplemented with 1 mM DTT, 1× PIC, and 1× PhosStop and incubated with anti-FLAG M2 agarose resin. Beads were washed, eluted in protein sample buffer, and analyzed using SDS-PAGE and Coomassie staining.

### Cellular fractionations

An approximately equal number of parental or endogenously tagged CDIN1-HF or CDAN1-HF Flp-In 293 T-REx cells were lysed using the NE-PER Nuclear and Cytoplasmic Extraction Kit (Thermo Scientific) supplemented with 1× PIC and 1× PhosStop. The cytosolic fraction was extracted following the manufacturer’s instructions. The remaining pellet containing the nuclei was washed and lysed in protein sample buffer with Benzonase (Millipore).

### Live-cell imaging and immunofluorescence

For live-cell imaging, cells were plated in either an 8-well chambered cover glass (Cellvis C8-1.5H-N) or a 12-well glass bottom plate. After 48 hr, cells were treated with 100 nM JFX650 HaloTag Ligand (a gift from A. Mizrak, the Harper lab, and the Lavis lab) for 1 hr at 37°C. Immediately before imaging, cells were stained with Hoechst (Invitrogen, H3570), washed, and placed in FluoroBrite DMEM (Gibco) supplemented with 10% FBS. Cells were mounted either in an OkoLab cage or stage top microscope incubator heated to 37°C with 5% CO_2_ for imaging.

For fixed imaging, cells grown on glass bottom plates were washed in PBS and fixed with 4% PFA in PBS for 10 min. Cells were then washed 3 times, permeabilized with 0.1% Triton for 5 min, then incubated with blocking buffer (10% FBS or 3% BSA in PBS with 0.05% Tween 20 [PBSt]) for 1 hr at room temperature. Primary antibody was diluted in blocking buffer and added at room temperature for 1 hr or 4°C overnight. After three PBSt washes, cells were incubated with Alexa Fluor 564-conjugated goat anti-rabbit IgG secondary antibody (1:500; Jackson ImmunoResearch, 111-585-003) in blocking buffer for 1 hr before labeling with Hoechst, then washed in PBSt.

All images were collected with a confocal Yokagawa CSU-X1 spinning disk confocal on a Nikon Ti inverted microscope equipped with either a Nikon Plan Fluor 40x Oil DIC H N2 (Supplemental Fig. 1d), Plan Apo 60× Oil DIC H objective (Fig. 1d), or Plan Apo λ 100x Oil objective (Supplemental Fig. 1e,f). Fluorescence was excited with solid-state lasers at 405 nm (80 mW), 561 nm (65 mW), or 640 nm (60 mW) and collected using ET455/50m, ET620/60m, or ET700/75m emission filters (Chroma), respectively. Images were acquired with a Hamamatsu ORCA-Fusion BT sCMOS camera controlled with NIS elements 5.21 software. Brightness and contrast were adjusted identically for compared image sets in Fiji^66^.

### Purification of CDAN1 complexes

ST-CDAN1 complexes were purified via co-expression with F-ASF1A (C:A) or both F-CDIN1 and HA-ASF1A (C:C:A), or via co-expression of truncated ST-CDAN1_C_ with F-CDIN1 (C:C), from Expi293 cells. Cells were seeded at 2.5 million cells/mL. 24 hr later, cells were diluted to 3 million cells/mL, and 0.5-1 μg/mL of each construct was transfected with 5 μg/mL polyethyleneimine (PEI-25K; Polysciences, 23966). 24 hr after transfection, 3 mM sodium valproate and 0.45% glucose were added to the cells. After 48 hr, the cells were collected, washed once in cold PBS, and either flash frozen and stored at -80°C for later processing or immediately lysed in 1 mL IP buffer supplemented with 1 mM DTT and 1× PIC per 10 mL culture. Lysates were clarified by centrifugation at 15,000-21,130×*g* for 10-20 min at 4°C. The supernatant was incubated with anti-FLAG resin for one hr at 4°C while rotating. The lysate was then passed through a column by gravity flow, washed with IP buffer, then with wash buffer (WB; IP buffer with 1 mM DTT but lacking Triton-X 100), and eluted using 3× FLAG peptide (APExBIO) in WB. The eluate was then incubated with Strep resin in WB for one hr at 4°C while rotating. After incubation, the sample was eluted using 10 mM desthiobiotin in WB.

### SEC-MALS

C:A and C:C:A complexes were purified as described. The elutions were pooled and applied to a Superose 6 10/300 GL column (Cytiva) pre-equilibrated in 1× PBS, pH 7.4 (Corning, 46-013-CM) with 1 mM DTT, and 0.2 mL fractions were collected. Peak fractions were pooled, concentrated, and 200 µg of each complex at 5 µM were applied to a pre-equilibrated SRT SEC-300 column (Sepax) connected to a DAWN HELEOS II Multi-Angle Light Scattering detector (Wyatt Technology) followed by an Optilab T-rEX Refractive Index Detector (Wyatt Technology) and a WyattQELS detector for Dynamic Light Scattering (Wyatt Technology). Data was analyzed in Astra 7 software (Wyatt Technology). Protein concentration was measured using refractive index; a refractive index increment (dn/dc) of 0.185 was used in MALS calculations. BSA was used as a standard (Thermo Fisher, 23209).

### Cryo-EM sample preparation and data collection

C:C:A complexes purified as described above were crosslinked with 250 µM BS3 (Thermo Fisher Scientific, 21580) for 15 min on ice and quenched with 2.5 mM Tris pH 7.6. 3 μL of the crosslinked sample at 1 mg/mL was applied to glow-discharged 0.6/1 UltrAuFoil 300 mesh grids (Quantifoil) and frozen in liquid ethane using a Vitrobot Mark IV (Thermo Fisher Scientific) set at 4°C and 100% humidity with a blot time of 3 seconds, a blot force of +8, and a wait time of 30s. CDAN1 C-term:CDIN1 (C:C) complexes were prepared as described above except crosslinked and plunge frozen at a final concentration of 0.78 mg/mL.

Datasets were collected using a Titan Krios (Thermo Fisher Scientific) operating at 300 kV and equipped with a BioQuantum imaging filter with a 20-eV slit width and a K3 direct electron detector (Gatan) in counting mode at a nominal magnification of ×105,000 corresponding to a calibrated pixel size of 0.825 Å. Semi-automated data collection was performed with SerialEM. For the C:C:A dataset, 2.708-second exposures were fractionated into 49 frames, resulting in a total exposure of 50.3 electrons/Å^2^. The defocus targets were −1.3 to −2.3 µm. For the C:C dataset, 2.597-second exposures were fractionated into 50 frames, resulting in a total exposure of 52.72 electrons/Å^2^. The defocus targets were −1.2 to −2.2 µm.

### Image processing and model building

Data processing was performed using cryoSPARC^67^ v4.3.1. Patch-based motion correction and CTF estimation were applied during cryoSPARC Live, and micrographs with severe contamination were removed. For the C:C:A complex, 3,250 micrographs were subjected to automated particle picking using templates generated from blob-based picking. Particles were initially extracted with a box size of 300 and downsampled to a box size of 128 for 2D classifications to remove junk particles. After *ab initio* reconstructions and heterogeneous refinements, 256,296 particles were unbinned and subjected to CTF refinement, local motion correction, and non-uniform and local refinements with C1 symmetry to produce a reconstruction of a CDAN1 dimer with two stacked ASF1A molecules at an overall resolution of 3.0 Å. The same particle set was also subjected to a non-uniform refinement with C2 symmetry that was used for symmetry expansion. Two rounds of 3D classification were performed with the expanded particle set with C1 symmetry using masks focused on the stacked ASF1A molecules associated with one CDAN1. This clearly demonstrated occupancy of one or two ASF1A molecules. Finally, 122,061 particles displaying occupancy for two ASF1A molecules were subjected to a final masked local refinement. To better visualize a CDAN1 dimer associated with 3 ASF1A molecules, a separate processing pipeline was employed using particles extracted in a larger box size of 500 and downsampled to a box size of 100 for initial ab initio refinement and 3D classification resulting in 25,075 particles with clear density for 3 ASF1A molecules that were unbinned and subjected to a final round of non-uniform refinement to produce a reconstruction at an overall resolution of 3.5 Å.

Half maps were post-processed using DeepEMhancer^68^ for interpretation. Alphafold2^69^ models of human CDAN1 (Q8IWY9) and ASF1A (Q9Y294) were used as initial models that were fitted as rigid bodies into the cryo-EM map of the C:C:A complex in ChimeraX^70^ v1.6. CDAN1 BD_A1_ and BD_A2_ placements were modeled using regions isolated from an Alphafold3^71^ prediction of one CDAN1 with two ASF1A molecules, aligned to the ASF1A models that were rigid body fitted into the map. The model was then manually adjusted in Coot^72^ v0.9 with multiple rounds of Phenix^73^ real space refine with manual adjustments in Coot in between each round. Refinements were first performed using a model with one copy of CDAN1 and two copies of ASF1A against the map focused on the two stacked ASF1A molecules, and then with the whole complex containing a CDAN1 dimer and three ASF1A molecules. Data processing was supported by software packages installed and configured by SBGrid^74^. Figure panels were made with ChimeraX. Initial multiple sequence alignments were generated using Clustal Omega^75^ and visualized using ESPript 3.0^76^. CDAN1 sequence conservation was analyzed using the ConSurf^63^ server.

## Notes

### Competing Interest Statement

The authors have declared no competing interest.

